# Temperature changes are signaled in cyanobacteria through the PipX interaction network

**DOI:** 10.1101/2025.05.19.654826

**Authors:** Antonio Llop, Sirine Bibak, Trinidad Mata-Balaguer, Lorena Tremiño, Laura Fuertes-García, José L. Neira, Ray Dixon, Asunción Contreras

## Abstract

Cyanobacteria perform oxygenic photosynthesis and have evolved sophisticated mechanisms to adapt their metabolism to challenging environmental changes. Despite their ecological and biotechnological importance, many regulatory proteins are still uncharacterised, and their signalling networks are poorly studied in comparison to other bacterial phyla. Two small proteins, PipX, unique to cyanobacteria, and PII, widespread in bacteria and plants, are the hubs of a protein interaction network involved in carbon/nitrogen homeostasis, energy sensing, translational regulation and growth. Here we exploit the NanoBiT complementation system to demonstrate in real time that temperature affects PipX interactions with its best studied partners: the signal transduction protein PII, the global transcriptional regulator NtcA, and the ribosome-assembly GTPase EngA. While heat shock increased PipX-PII complex formation and impaired PipX-EngA and PipX-NtcA interactions, cold shock resulted in a decrease of all three complexes. Far-UV circular dichroism spectra of isolated PipX suggested the involvement of its C-terminal α-helix in the common response to cold shock. However, during longer term acclimatization, each type of complex responded distinctively after up- or downshifts in temperature and PipX-PII and PipX-NtcA interactions were influenced in opposite ways. Altogether the results indicate that PipX is a thermometer of low temperatures, bringing new light to the study of environmental signaling in cyanobacteria. Our results also illustrate the enormous potential of the NanoBiT complementation system to fuel understanding of the mechanisms allowing cyanobacteria to initially respond and/or acclimatize to environmental factors.

**IMPORTANCE:** Cyanobacteria are a group of organisms of great ecological and biotechnological importance but relatively little understood in terms of the regulatory components and molecular mechanisms that make them so unique. PipX is a small protein exclusive to cyanobacteria that functions by binding to other regulators in response to intracellular metabolic signals. We used a bioluminescence reporter system to show that temperature shifts significantly alter the relative affinity of PipX for its well-known partners. By showing the impact of a highly relevant environmentally factor such as temperature on the regulatory details of a protein interaction network and implicating PipX in the response to cold shock this work paves the way for significant advancements in both basic and applied research of cyanobacteria.

## INTRODUCTION

Cyanobacteria, phototrophic organisms that perform oxygenic photosynthesis, constitute an ecologically important phylum that is responsible for the evolution of the oxygenic atmosphere. They are the main contributors to marine primary production (1, 2) and are also ideal production systems for several high-value compounds, including biofuels (3). Cyanobacteria have developed sophisticated regulatory systems to adapt to challenging environmental conditions, including strategies to maintain homeostasis of the carbon/nitrogen balance (reviewed by (4, 5)).To achieve this homeostasis the signal transduction protein PII regulates the activity of proteins involved in nitrogen and carbon metabolism by direct protein-protein interactions (6), perceiving metabolic information through the competitive binding of ATP or ADP and the synergistic binding of ATP and 2-oxoglutarate (2-OG) (7, 8). The global transcriptional regulator NtcA controls nitrogen assimilation in cyanobacteria (9–11) by also responding to the concentration of 2-OG, which provides a metabolic sensor of the carbon and nitrogen status.

The PipX protein (12), identified by its ability to form complexes with PII and NtcA (9, 13–18), is a unique protein exclusive to cyanobacteria. Regulation of protein-protein interactions with PipX is dependent on ligand binding by its partners (Fig. 1). PII stabilizes PipX in *Synechococcus elongatus* PCC 7942 (hereafter *S. elongatus*) (19). The binding of PipX to PII or NtcA is antagonistically tuned by 2-OG levels (9, 16, 20). PipX uses the same surface from its TLD (Tudor-like domain) /KOW domain to bind to either 2-OG-bound NtcA, stimulating DNA binding and transcriptional activity, or to 2-OG-free PII; thus PII sequestration of PipX at low 2-OG reduces the expression of NtcA-dependent gene targets (21–26). PipX stabilizes the conformation of NtcA that is transcriptionally active and helps local recruitment of RNA polymerase (27) in response to nitrogen limitation (24, 28). PipX also interacts with the essential ribosome-assembly GTPase EngA (YphC/Der/YfgK) (29). In *S. elongatus*, PipX interferes with EngA function under environmentally relevant conditions such as cold or light stress (29, 30).

**Figure 1.**
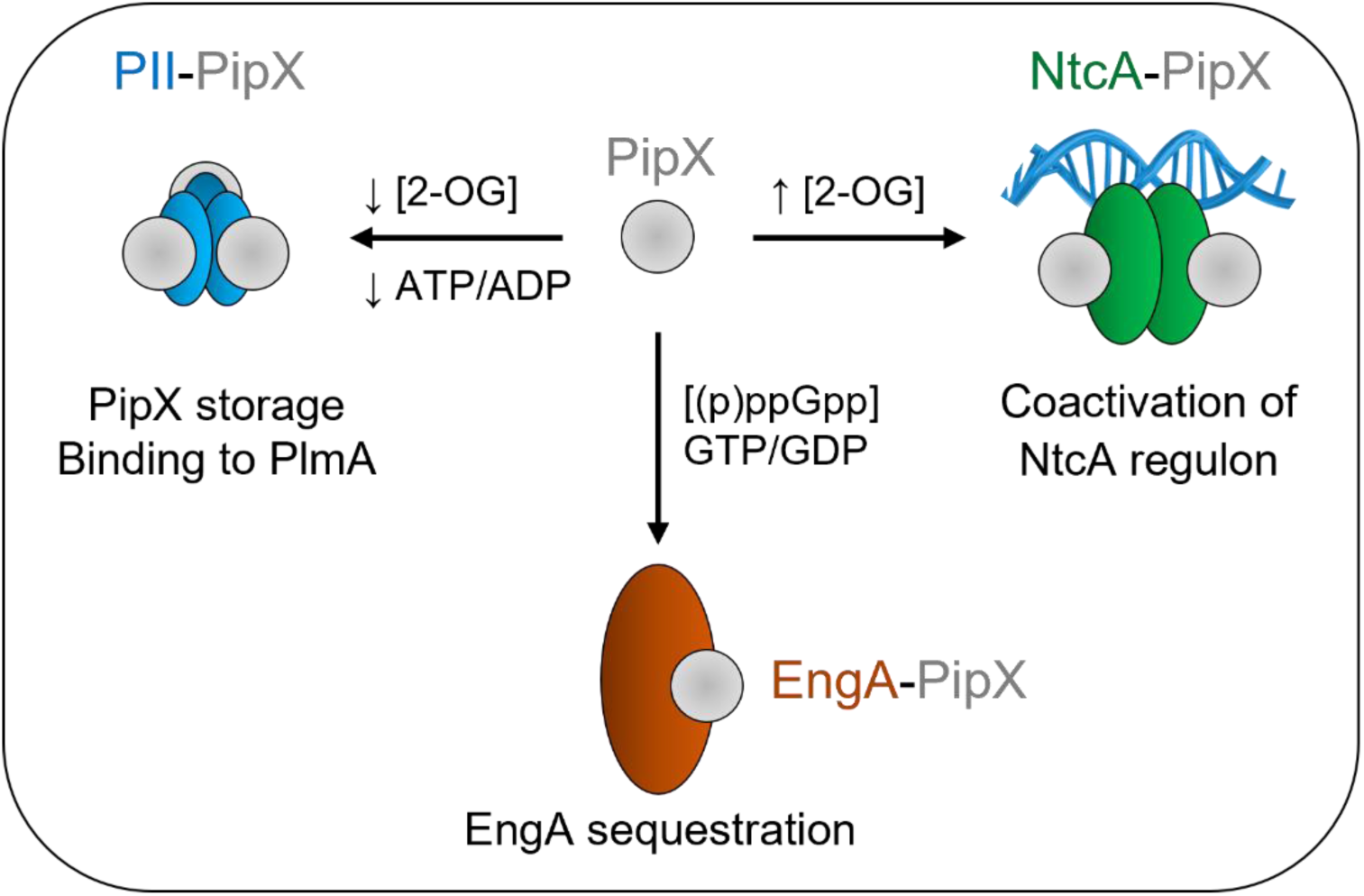
Functions and effectors of PipX complexes. Geometric representation of the indicated proteins in their corresponding oligomeric state, with area scaled according to the number of amino acids. Known or putative effectors involved complex formation and their functions are indicated. See text for additional details.

Temperature is a highly important environmental parameter for cyanobacteria with a considerable impact on both physiology and gene expression. Previous studies have focused on identifying regulators and gene targets of signal pathways involved in transcriptional regulation in response to either cold or heat shock (31–33), two stresses that are sensed by different mechanisms in bacteria (34–39). An alternative approach, studying the in vivo effect of temperature shifts on regulatory protein complexes is now possible using the NanoBiT complementation system (40), which is based on reconstitution of the small and high output bioluminescence enzyme NanoLuc. The NanoBiT system has been used in both mammalian (41–43) and bacterial cells (18, 44–48) to demonstrate the specificity of protein interactions of interest in their natural environment. Importantly, we have used it to demonstrate the opposing regulation of PipX-PII and PipX-NtcA complexes in real time in response to different nitrogen sources or to decreases in ATP levels in *S. elongatus* (18). These studies highlight the advantages of the NanoBit system in determining real time effects of ligands on complex formation and the competition for PipX between the two nitrogen regulators under environmentally relevant conditions for cyanobacteria.

Since temperature appears to be highly relevant for EngA function and interactions with PipX (29, 30), we have used the NanoBit system to investigate the importance of temperature on the PipX interaction network in vivo. We demonstrate that temperature regulates not only EngA levels and PipX-EngA complex formation but also the stability of PipX-PII and PipX-NtcA complexes in *S. elongatus*. Real-time experiments using the NanoBiT complementation system revealed the involvement of PipX in both early and late responses. The rapid decrease upon cold shock of PipX affinity for all three partners, in combination with far-ultraviolet (UV) circular dichroism (CD) experiments with purified PipX, suggested that temperature downshift induce conformational changes in PipX. This study illustrates the enormous potential of the NanoBiT complementation system to fuel our understanding of molecular mechanisms allowing cyanobacteria to initially respond and acclimatize to environmental conditions.

## RESULTS

### EngA levels were downregulated by high temperature at the post-transcriptional level

We recently showed that the levels of the ribosome-assembly GTPase, EngA, increase after transfer from 30°C to 18°C while PipX or PII levels remain constant (30). To investigate whether temperature upshift also affected EngA levels, we determined EngA, PipX and PII levels at different timepoints after transferring cultures from 30°C to 42°C. Western blot analysis with anti-EngA or anti-PipX antibodies were performed at different timepoints after transferring *S. elongatus* cultures from 30°C to 42°C. To quantify the changes and compare the signal intensity for EngA or PipX bands, the immunodetection signal for PlmA was used as an internal control to normalize signals and determine protein levels at 42°C for each protein. As an additional control, we determined PII levels in parallel, showing that they were also indistinguishable among the different samples. As shown in Fig. 2*A* and S1*A*, EngA levels were significantly lower at 42°C, decreasing to less than 50% of the 30°C level. The significant decrease in EngA levels, detected 3 hours after the upshift, was maintained for 24-hours, with no significant changes between the time points. In contrast, PipX levels remained constant after the temperature upshift.

**Figure 2.**
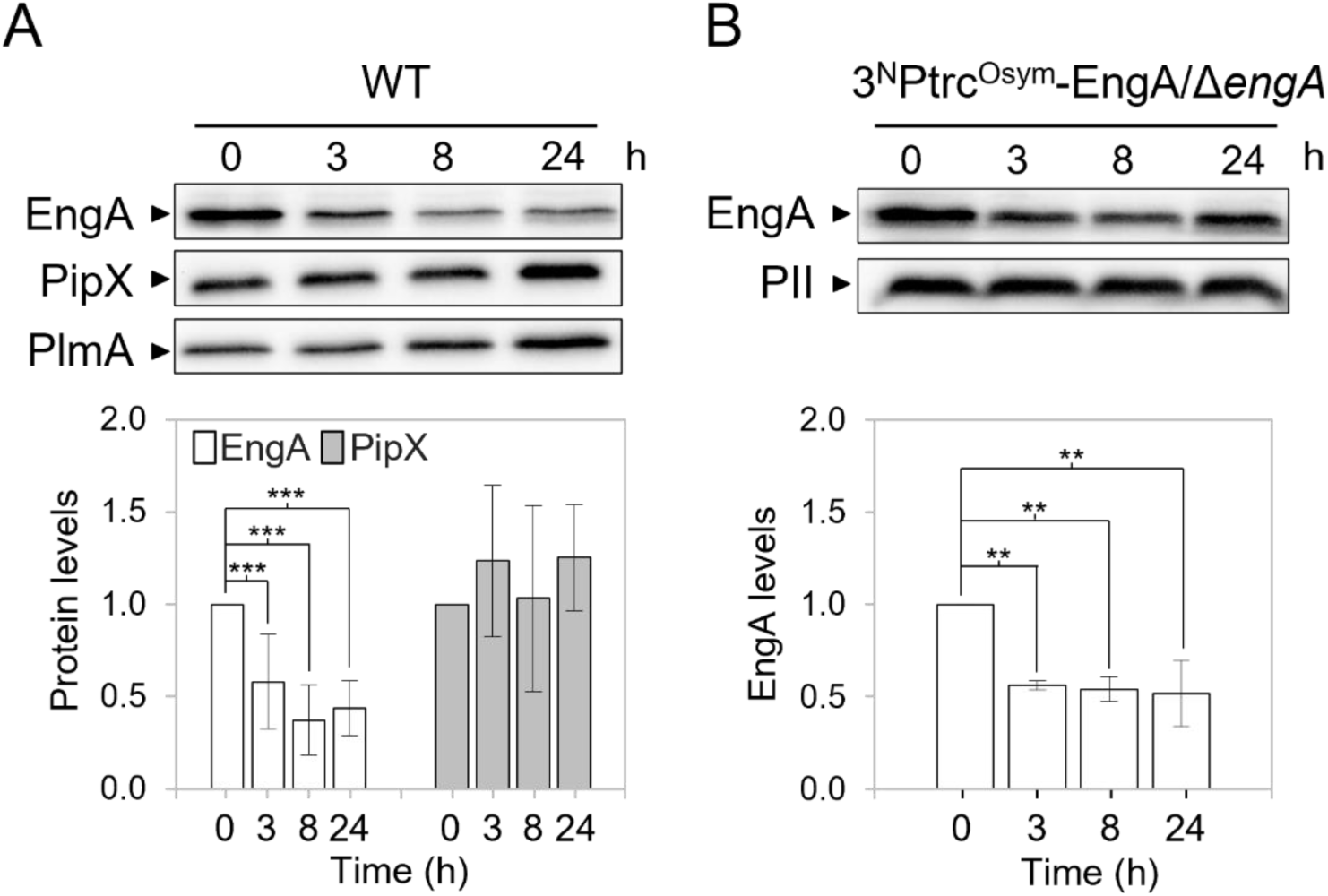
Effect of the temperature upshift on protein levels in *S. elongatus*. (*A, B*) *Top* – Representative immunodetections of the indicated proteins in the WT (*A*) and 3^N^Ptrc^Osym^-EngA/*engA* (*B*) strains at 42°C. *Bottom* – Relative protein levels normalized to PlmA (*A*) or PII (*B*) and referred to the timepoint 0. Data are presented as means with error bars (standard deviation) from at least four biological replicates. A linear mixed model was performed with time as a fixed effect and experiment as a random effect. Comparison of protein levels between timepoint 0 and the others were made using pairwise comparisons with Kenward-Roger adjusted degrees of freedom and Bonferroni correction. Significance levels were denoted as p ≤ 0.01 (**) and p ≤ 0.001 (***).

To gain insights into the mechanism involved in down regulation of EngA levels at 42°C, we next generated the *S. elongatus* strain 3^N^Ptrc^Osym^-EngA*/*Δ*engA* (Table 1), where expression of the ectopic *engA* gene (allele *Ptrc^Osym^*::*engA*) is driven from an IPTG inducible promoter while coding sequences at the native *engA* locus were precisely replaced by the *cat* (chloramphenicol-acetyltransferase) gene (allele *engA*::*cat*). Cultures of this *S. elongatus* strain were transferred from 30°C to 42°C and extracts taken at different timepoints were subsequently analyzed by Western blotting. As shown in Fig. 2*B* and S1*B*, EngA levels decreased in strain 3^N^Ptrc-EngA*/*Δ*engA* at 42°C, indicating that *cis*-acting sequences upstream of the *engA* gene were not required for downregulation in response to high temperature and thus decreased translation and/or increased degradation of EngA occurred at 42°C.

**Table 1.**
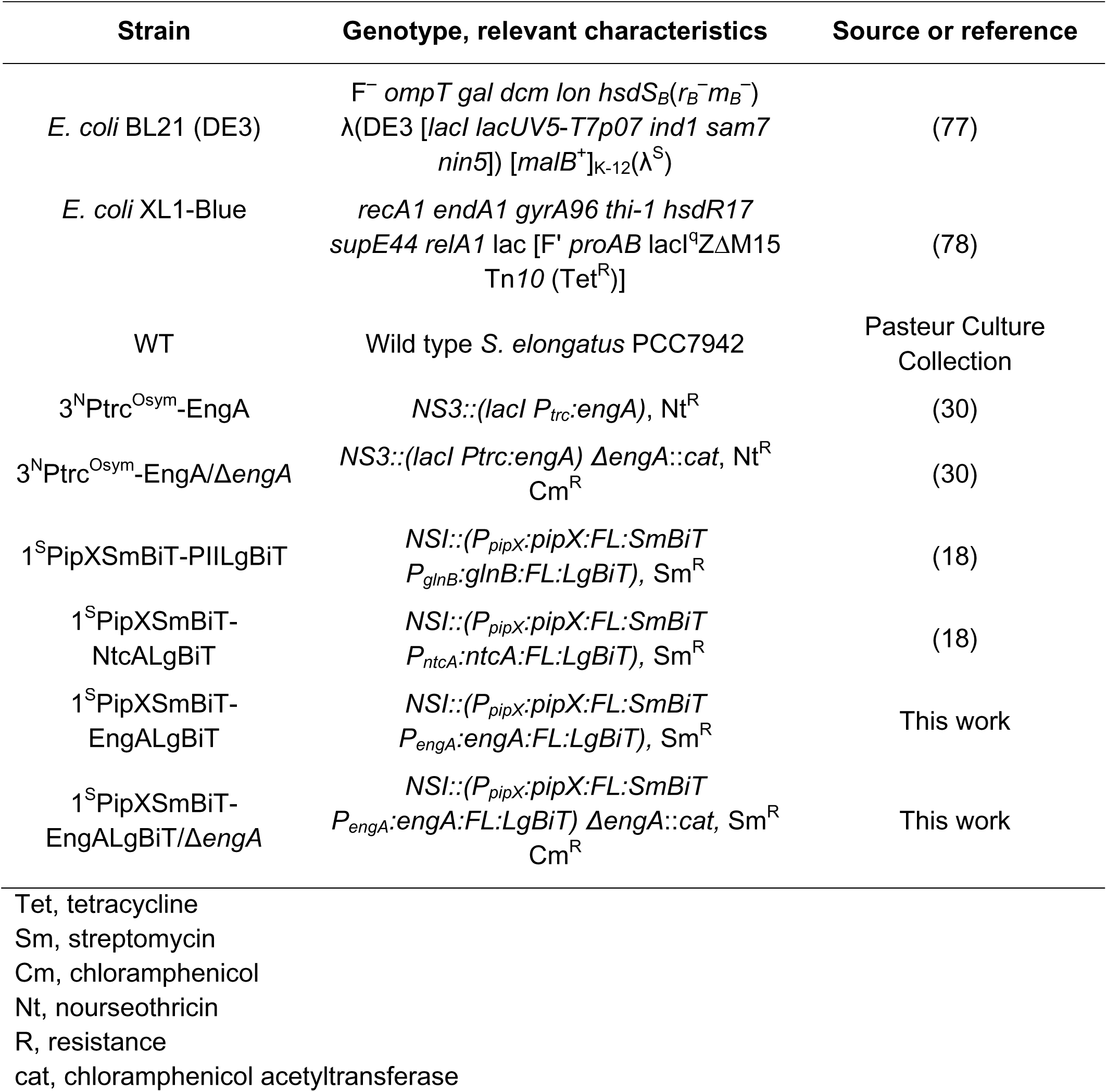
Strains.

In combination with our previous experiments performed at 18°C (30) it is evident that overall, EngA levels decreased in *S. elongatus* in response to temperature transitions to warmer environmental conditions, consistent with the role of EngA as an rRNA chaperone at low temperatures (49). However, it is evident that the mechanistic response to temperature stress was different, since cold shock induced transcriptional regulation of EngA expression, whereas the heat shock response involves post-transcriptional control of EngA levels. This raises the question of whether the levels of other proteins acting as RNA chaperones were also finely tuned in response to temperature in cyanobacteria.

### Generation of a NanoBiT reporter strain for PipX-EngA interactions in *S. elongatus*

To determine the effect of temperature on PipX-EngA complex formation in *S. elongatus* we designed a PipX-EngA reporter construct guided by previously validated PipX-PII and PipX-NtcA reporters (18). This construct expresses PipX-SmBiT and EngA-LgBiT fusion proteins from a neutral site in the *S. elongatus* chromosome (Fig. 3*A*). To conserve wild type regulation, the upstream regulatory sequences of *pipX* and *engA* were also included. Introduction of the PipX-EngA reporter construct into the neutral site I (NSI) by allelic replacement was facilitated by a streptomycin-resistant marker cassette (C.S3) also included within the NSI insertions.

**Figure 3.**
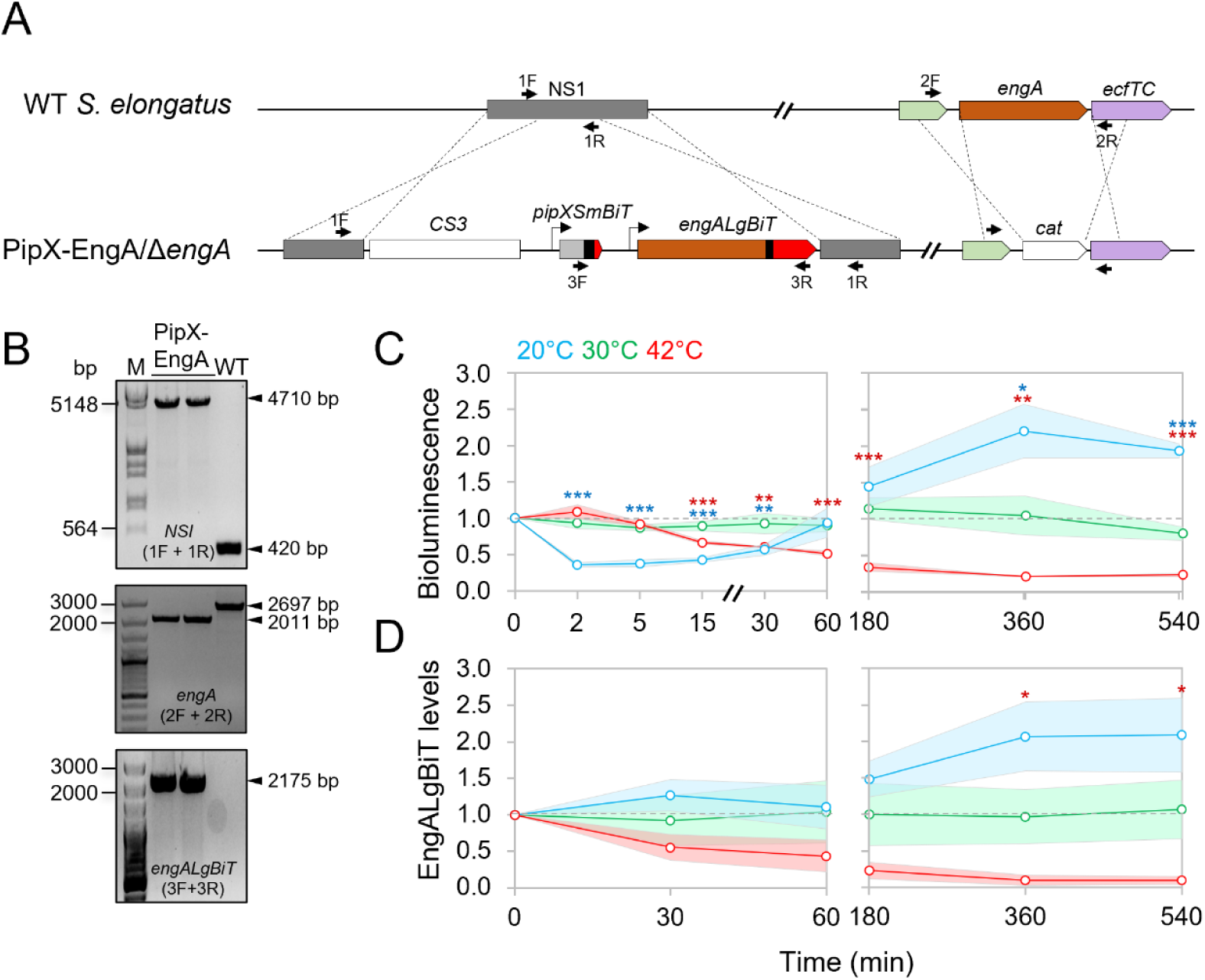
Construction and validation of a PipX-EngA NanoBit reporter and its response to temperature shifts in *S. elongatus.* (*A*) Schematic representation of NSI and *engA* regions at the *S. elongatus* chromosome from WT (top) or reporter strain (botton) showing *pipXSmBiT* and *engALgBiT* gene fusions and *CS3* and *cat* selection markers. The color code for *pipX* and *engA* genes is the same used for the proteins in Fig.1. Black bars indicate flexible linkers. (*B*) PCR analysis with the primers indicated as black arrows in *A*. M: λ HindIII/EcoRI or 100 bp size marker. (*C*) Bioluminescence signals obtained using Hikarazine-108 (CNRS) as the luciferase pro-substrate. Signals were referred to the timepoint 0, in cultures at 20°C, 30°C, or 42°C corresponding to timepoints taken up to one (*left*) or nine hours (*right*). The time axis has been interrupted to improve data visualization. (*D*) EngALgBiT levels, normalized to PII and referred to timepoint 0, corresponding to timepoints taken up to one (*left*) or nine hours (*right*). Dashed grey lines mark the threshold at 1 in the graphs. Data are presented as means with error bars (standard deviation as shadows behind the lines) from at least three biological replicates. Welch’s t-test with Bonferroni correction was used to compare data between 30°C and either 20°C or 42°C at the same timepoint. Significance levels were denoted for the corresponding color of the condition as p ≤ 0.05 (*), p ≤ 0.01 (**), and p ≤ 0.001 (***).

The PipX-EngA reporter plasmid (pUAGC1165, Table 2), was introduced into *S. elongatus* and independent streptomycin-resistant transformants were PCR-analysed to confirm complete segregation of the modified NSI alleles in *S. elongatus*. (named NSI in Fig. 3*B*). Validated clones were selected for further work (strain 1^S^PipXSmBiT-EngALgBiT; Fig. 3*B*, upper panel).

**Table 2.**
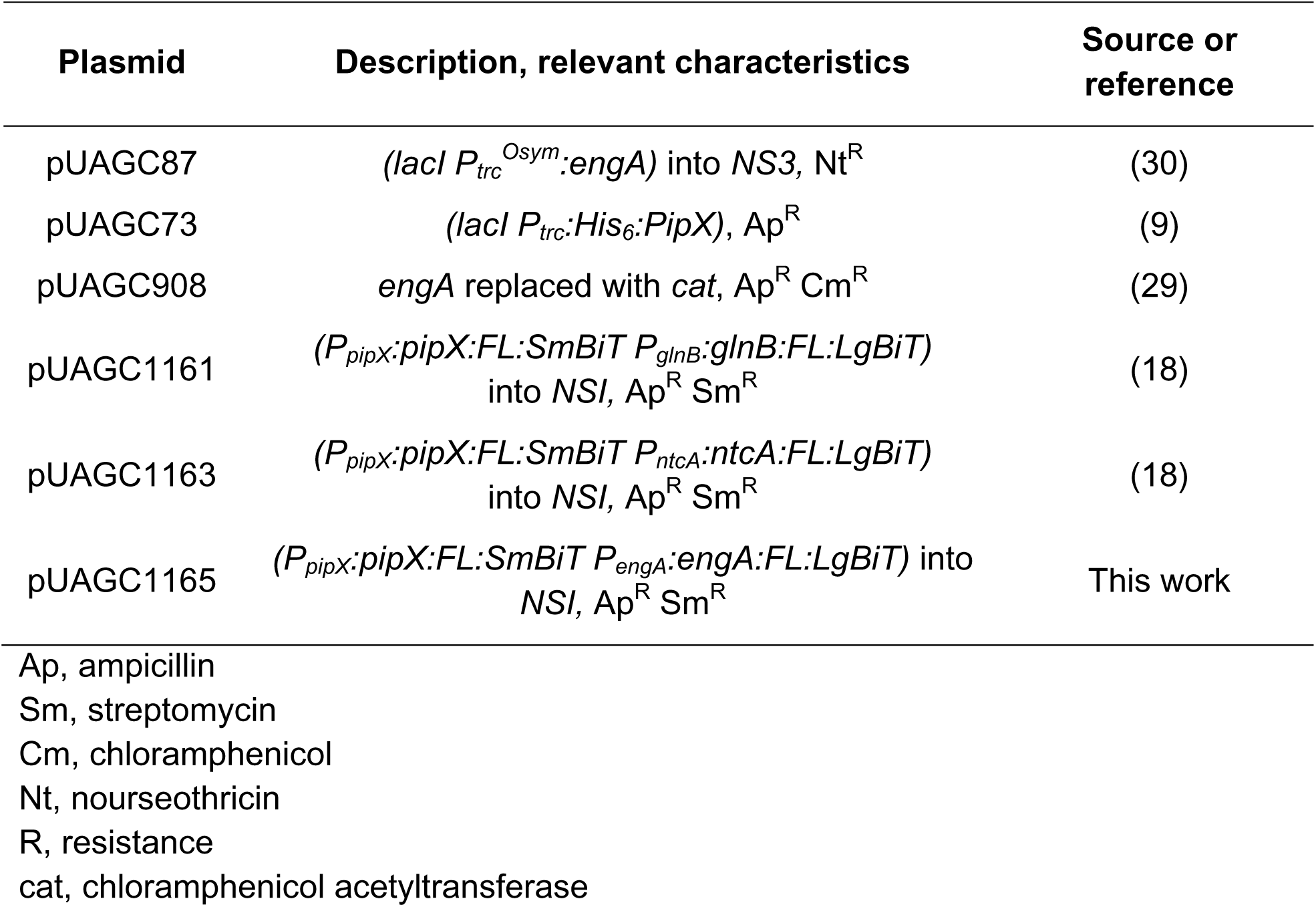
Plasmids.

| Plasmid | Description, relevant characteristics | Source or reference |
| --- | --- | --- |
| pUAGC87 | (*lacI* $P_{trc}^{Osym}$:*engA*) into NS3, Nt<sup>R</sup> | (30) |
| pUAGC73 | (*lacI* $P_{trc}$:*His*<sub>6</sub>:*PipX*), Ap<sup>R</sup> | (9) |
| pUAGC908 | *engA* replaced with *cat*, Ap<sup>R</sup> Cm<sup>R</sup> | (29) |
| pUAGC1161 | ($P_{pipX}$:*pipX*:*FL*:*SmBiT* $P_{glnB}$:*glnB*:*FL*:*LgBiT*) into NSI, Ap<sup>R</sup> Sm<sup>R</sup> | (18) |
| pUAGC1163 | ($P_{pipX}$:*pipX*:*FL*:*SmBiT* $P_{ntcA}$:*ntcA*:*FL*:*LgBiT*) into NSI, Ap<sup>R</sup> Sm<sup>R</sup> | (18) |
| pUAGC1165 | ($P_{pipX}$:*pipX*:*FL*:*SmBiT* $P_{engA}$:*engA*:*FL*:*LgBiT*) into NSI, Ap<sup>R</sup> Sm<sup>R</sup> | This work |

Ap, ampicillin Sm, streptomycin Cm, chloramphenicol Nt, nourseothricin R, resistance *cat*, chloramphenicol acetyltransferase

Since *engA* is essential in *S. elongatus*, we next tested the functionality of the EngA-LgBiT fusion protein by its ability to provide EngA essential functions. To this end, *S. elongatus* strain 1^S^PipXSmBiT-EngALgBiT was used to inactivate the native *engA* gene by allelic replacement with the *engA::cat* derivative. The corresponding alleles are illustrated in Fig. 3*A*. Complete segregation of the null allele indicated complementation of EngA essential functions by EngA-LgBiT. Validated 1^S^PipXSmBiT-EngALgBiT *engA* clones expressing *engALgBiT*, instead of the wild type *engA* allele (Fig. 3*B*, middle and lower panels), were selected and referred to hereafter as the PipX-EngA reporter strain.

### PipX-EngA complexes were transiently impaired by cold shock, but increased in cultures acclimated to low temperature

To test the effect of temperature changes on PipX-EngA complexes in real-time we measured bioluminescence at different timepoints after shifting cultures of the corresponding NanoBiT reporter strain from 30°C to 20°C or 42°C. Luminescence values were also recorded from control samples maintained at 30°C. To facilitate the distinction between the initial responses to cold or heat stress from longer term acclimatization responses, we split data from the same experiments into independent figures to represent “early” (up to 60 min, Fig. 3*C* left) and “late” (up to 9 hours, Fig. 3*C* right) timepoints. Given that at any given timepoint bioluminescence signals from NanoBit reporters were affected by both the levels and the affinity between the two partners, and EngA levels were temperature-dependent in wild type *S. elongatus* ((30); Fig. 2), samples were taken at several of the timepoints used for luciferase assays and subsequently analysed by Western blot with anti-LgBiT (Fig. 3*D*, S1*C* and S2*A*). Transfer to 20°C produced an extremely rapid decay of the bioluminescence signal, (by about 70% at the 2 min timepoint) indicating that cold shock dissociated PipX-EngA complexes. In contrast, transfer to 42°C resulted in very slow decrease of the PipX-EngA signal that mirrored the decrease in EngALgBiT levels. The fast decay of the bioluminescence signal upon transfer to 20°C was fully recovered at the 60 min timepoint, with signals continuing to increase and reaching maximal values at the longest timepoints of the experiment, where they closely correlated with EngALgBiT levels (Fig. 3*C* and *D*). This implies that the immediate response to cold shock resulted in complex dissociation, whereas acclimatization allowed reestablishment of complexes in the longer term, in agreement with the increase in EngA levels. In contrast, close correlation between bioluminescence signals and EngALgBiT levels were observed between all timepoints taken from cultures at 42°C, indicating that there was no rapid response of complexes to heat shock and that PipX-EngA complexes were maintained at this temperature, even though there was a significant decrease in EngA levels.

Considering the relative bioluminescence values under steady-state conditions (longer term exposure to temperature) a larger amount of PipX-EngA complexes were formed in cultures exposed to cold rather than to heat stress, raising questions concerning the function of PipX-EngA complexes in cells acclimatized to low temperature.

### PipX-PII and PipX-NtcA complexes were rapidly impaired by cold shock, but responded differently to heat shock

PipX-PII and PipX-NtcA complexes have been extensively studied as part of a protein network for metabolic regulation and signaling the intracellular carbon/nitrogen and energy status, but so far not in other contexts that may also be relevant to the PipX interaction network. Similar to the analysis of PipX-EngA complexes, we utilized the Nanobit system to determine the effect of temperature changes on PipX-PII and PipX-NtcA complexes at different timepoints after shifting cultures from 30°C to 20°C or 42°C (Fig. 4*A*). Western blot analyses with anti-LgBiT indicated that the levels of both PIILgBiT and NtcALgBiT remained relatively constant during the time course of the experiment (Fig. 4*B*, S1*D*, and S2*B* and *C*). Thus, differences in the bioluminescence signal after the temperature shifts should directly report the in vivo dynamics of PipX-PII and PipX-NtcA complexes in response to cold or heat shock. Changes in bioluminescence signals took place rapidly (within 2 min) after each of the two temperature shifts (Fig. 4*A*). As in the case of the PipX-EngA interaction (Fig. 3*C*), cold shock decreased bioluminescence signals from both PipX-PII and PipX-NtcA reporters, indicating that a common mechanism could be involved in signaling cold shock to all three PipX complexes studied in this work. In view of this uniform response, it is unlikely that the changes in known ligands (ATP/ADP ratio or the 2-OG concentration), that influence partner switching of PipX between PII and NtcA (Fig. 1) were involved. Moreover, we detected no significant changes in the ATP levels during the time course after switching from 30°C to 20°C (Table S1).

**Figure 4.**
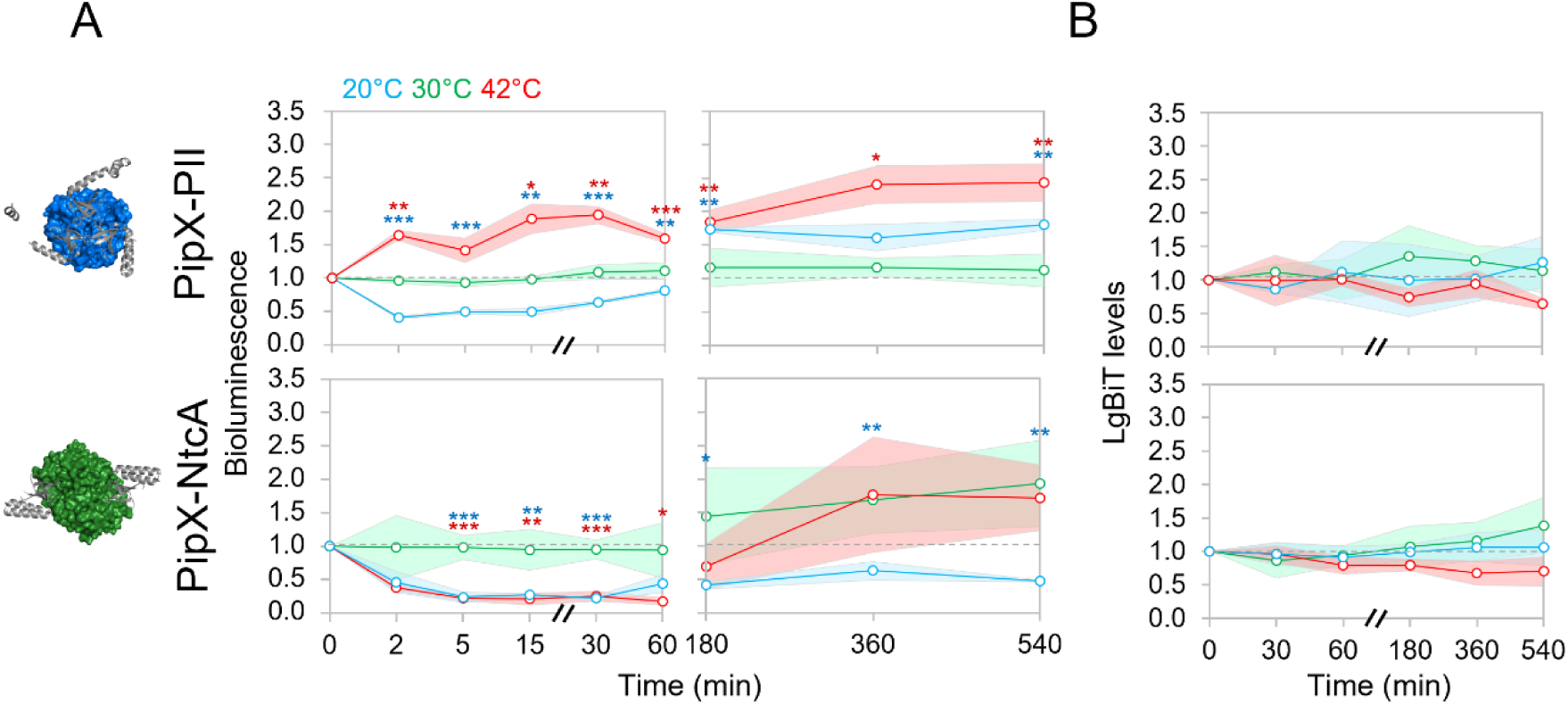
Regulation of PipX complexes in response to temperature up or downshifts. (*A*) Bioluminescence signal, referred to the timepoint 0, in *S. elongatus* cultures at 20°C, 30°C, or 42°C corresponding to timepoints taken up to 1(*left*) or nine hours (*right*). (*B*) Protein levels normalized to PlmA (for PIILgBiT) or PII (for NtcALgBiT) and referred to the timepoint 0. PipX (grey; ribbon) in complex with PII (blue; surface; PDB: 2XG8) or NtcA (green; surface; PDB: 2XKO) is shown at the left. Other details as in Fig 3.

In contrast, heat shock triggered opposite responses for PipX-PII and PipX-NtcA complexes, respectively increasing or decreasing complex formation, reminiscent of partner switching. Although we found no significant differences between the intracellular levels of ATP from *S. elongatus* cultures at 30°C or 42°C (Table S1), 2-OG levels were not investigated, and the results were still in agreement with the involvement of metabolic effector ligands in signaling.

### PipX-PII complexes were abundant during adaptation to both cold and heat stress

At 20°C the initial cold shock-promoted dissociation of PII-PipX complexes slowly reversed and between the 3h and 9h timepoints, bioluminescence signals were maintained above the control levels at 30°C, while PipX-NtcA signals remained minimal (Fig. 4*A*). The contrasting behavior of PipX-PII and PipX-NtcA complexes after acclimatization to low temperatures suggests that binding of PipX to PII was favored to the detriment of PipX-NtcA complexes. It is therefore likely that the NtcA regulon was down regulated as part of the acclimatization response to low temperature stress.

The initial increase in PipX-PII complex formation at 42°C remained above the control levels at 30°C during the 9 hours time course suggesting a role for PipX-PII complexes in acclimatization to heat stress. On the other hand, bioluminescence signals from the PipX-NtcA reporter were highly variable towards the end of the experiment at both 42°C and 30°C and thus, they were not easily rationalized. However, the signals from the PipX-NtcA complexes at 20°C were more consistent during the later stages of the time course, congruent with a model in which modulation of the NtcA regulon is necessary for temperature adaptation as noted above.

### Temperature induced conformational changes in PipX structure

The finding that shifting cultures from 30°C to 20°C impaired PipX complexes with PII, NtcA and EngA implied a sudden decrease in the affinity of PipX for all three partners, prompting us to explore the possibility that lowering the temperature affected somehow PipX conformation.

To investigate the influence of temperature on the conformation of PipX, we used far-ultraviolet circular dichroism (far-UV CD) to probe the conformation of recombinant PipX at temperatures in the range 10°C to 40°C, well below the thermal denaturation midpoint of the protein (61°C, as measured by thermal denaturation in the far-UV CD spectrum and by fluorescence emission after excitation at 280 nm.; Fig. S3). We observed that at 222 nm (the wavelength where α-helices exhibit one of the two minima spectral signatures (50)), there was a change in the variation of the ellipticity, which decreased almost linearly from 10°C to 25°C (Fig. 5*A* and *B*); but above 25°C the ellipticity reached a plateau, with smaller variations as the temperature further increased (Fig. 5*B*).

**Figure 5.**
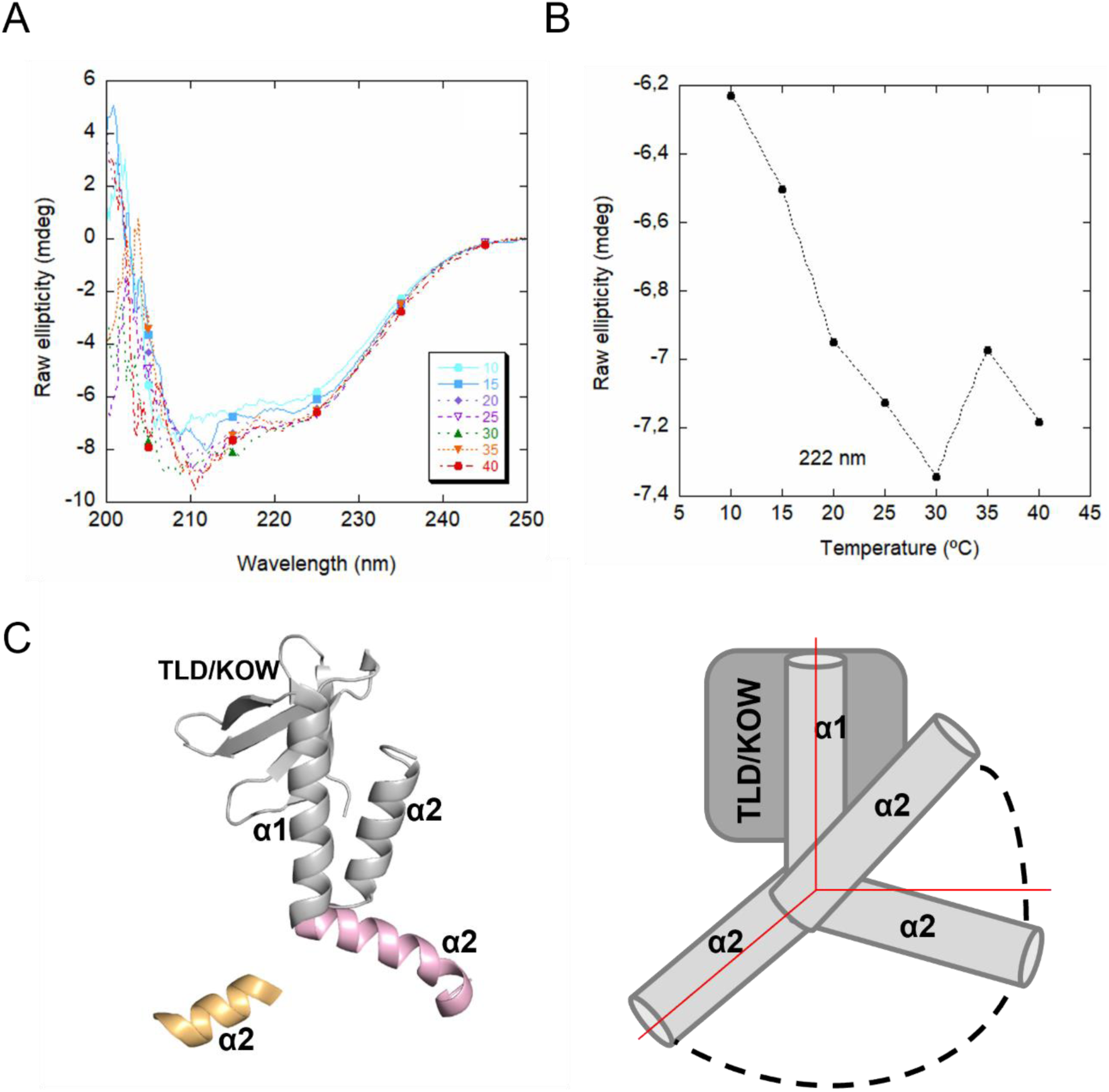
Temperature-dependent changes of the PipX secondary structure as monitored by far-UV CD. (*A*) Far-UV CD steady state spectra of PipX at different temperatures at 50 mM Tris pH 7.5. (*B*) Variations at one selected wavelength of the ellipticity (from the spectra in *A*) as the temperature was modified. (*C*) *Left* – The flexed form of PipX (grey) with alternative conformations of PipX α2 helix (pink and orange, corresponding to the extended, flexed and untraced forms) superimposed on it, as detected in the PipX-PII crystal structure complex (PDB: 2XG8). *Right* – Model of PipX α2 helix movement based on PipX crystal structures. Red lines represent spatial axes.

The structure of PipX, when bound to NtcA or PII, consists of an N-terminal, five-stranded β-sheet (comprising the Tudor-like or KOW domain), and two long C-terminal α-helices (16, 20). These helices adopt either: (i) a flexed (or closed) conformation in the PipX-NtcA and PipX-PII complexes, where they are in close contact in an antiparallel orientation, packing against the β-sheet, as in the isolated PipX protein (51); or, alternatively (ii) an open (or extended) conformation, observed in two of the three PipX molecules in the PipX-PII complex, where the last α-helix moves apart from the rest of the protein (Figs. 4*A* and 5*C* left) (16, 20). While NMR studies of isolated PipX in solution revealed that the C-terminal α-helices adopt the flexed conformation at 17°C (51), the inherent flexibility of the last α-helix is likely to allow additional conformations of the C-terminal α-helical domain, as suggested by the changes in the far-UV CD spectra at higher temperatures.

In the light of the data presented here, we propose that shifting cultures from 30°C to 20°C could displace the equilibrium for the different conformations of PipX (or intermediate ones) that have been detected in the crystal structures of the complexes or in isolation (Fig. 5*C* right), favoring conformations that reduced the affinity for PII, NtcA and EngA, and potentially increased it for unknown PipX partners.

## DISCUSSION

Temperature is a very important environmental parameter for most biological processes that directly impacts the structure and function of different cellular components with thermodynamically sensitive proteins. Responses to temperature shifts have been widely investigated in cyanobacteria and other bacterial groups. So far, the main experimental approaches to unravel the molecular details involved have focused on identifying regulatory proteins triggering a transcriptional response and on their gene targets. The former includes two-component systems where a sensor histidine kinase detects membrane fluidity and transmits the signal to its cognate partners by phosphorylation (31–36, 38, 39), although additional complexity is also emerging from those studies (52). Protein folding is also a process expected to be hindered by low temperature, yet its effect on cold shock response is poorly studied (36). In this work, we used a very different approach to gain insights into the mechanisms by which temperature can be sensed and affect signaling pathways. Taking advantage of the NanoBit complementation system, we report here on the effects, in real time, of temperature shifts on protein complexes belonging to the paradigmatic PipX interaction network of cyanobacteria.

We have shown in this work that the PipX interaction network is highly influenced by temperature, which could act at different regulatory levels to alter the distribution of PipX into different protein complexes. In the case of EngA, adaptation to temperature resulted in significant changes in the signal for PipX-EngA complexes, which were much more abundant at low than at high temperatures, reflecting the temperature-dependence of EngA levels. However, this was not an issue for PipX-PII or PipX-NtcA complexes, which exhibited adaptations to temperature during acclimatization and here the effect was likely to be attributed to altered levels of the ligands affecting complex formation due to the metabolic changes triggered by different culture temperatures. In contrast to the long-term adaptation to temperature, the response of PipX-PII, PipX-NtcA and PipX-EngA complexes to heat or cold shock was rapid and transient, corresponding to fast changes in affinity that although equilibrate towards a steady state during adaptation to temperature stress, cannot be attributed to changes in the levels of the interacting proteins. Importantly, the significant impairment of PipX-PII, PipX-NtcA and PipX-EngA complexes after cold shock cannot be easily explained based on the known effectors of these proteins.

It is worth noting that PII and PII-like proteins can bind and be regulated by different types of nucleotides including the second messengers cyclic adenosine monophosphate (cAMP) and cyclic di-adenosine monophosphate (3’,5’-cAMP) (53), including, (p)ppGpp, cyclic di-GMP and 2’,3’-cAMP have been shown to be involved in signalling cold shock in Gram positive bacteria or plants (54, 55). EngA proteins also bind (p)ppGpp (56–59). Unfortunately, cold is not one of the stress conditions that have been investigated in connection with the stringent response and (p)ppGpp levels in cyanobacteria (60–62), and we cannot safely rule out the involvement of additional ligands at this stage. However, the rapid generic dissociation of PipX from its partners observed upon shift to low temperature, in addition to the conformational changes at the C-terminal domain of PipX in response to temperature suggested by far-UV CD spectra, support a role for PipX as a direct sensor of cold shock. The movement of the C-terminal α-helix, as a consequence of its inherent flexibility during different time-scales (51), could modulate the binding of PipX to its partners. It is thus tempting to propose that temperature modulates the equilibrium towards different orientations of the C-terminal α-helix without altering the overall scaffolding of PipX and that this α-helix could act as a gate-keeper, allowing accessibility towards the TLD/KOW domain and the rest of PipX.

The common response to cold shock of all three complexes studied here raises questions regarding the existence and regulatory relevance of a wider PipX interaction network, that may involve binding partners that are favored immediately after the temperature shift. In this context, we favour the idea that cold shock triggers the sequestration of PipX by cellular components that form part of abundant protein or riboprotein complexes. This hypothesis is fuelled by information from several pieces of evidence. First, PipX appears to act as a growth brake under stress conditions or when overexpressed, inhibiting diverse relevant processes including photosynthesis (29, 30, 62, 63) and EngA-dependent ribosome assembly/translation during cold or high light stress (29, 30). Secondly, it colocalizes with the RNA-protein complexes involved in transcription, RNA metabolism and transcription initiation (64). Finally, it can interact with the sigma and gamma subunits of the RNA polymerase (27). Whatever the case, the present results provide clues to the identification of additional components of the PipX interaction network.

The finding that temperature shifts affected the relative abundance of PipX partner interactions is not very surprising in the light of the importance of temperature for gene expression and cell physiology. Changes in the PipX interactions observed after several minutes to hours would mainly impact metabolism, gene expression and ribosome assembly acting via PII, NtcA and EngA, respectively. However, decreased PipX interactions of PipX with those partners upon a sudden drop of temperature is likely to favor the binding of PipX to other, yet unknown, partners. The main questions now are the identity of these hypothetical partners and the physiological consequences of the transient PipX complexes inferred in this work.

Temperature downshift decreases photosynthesis (65, 66) and triggers photodamage because the rate of electron consumption decreases while the light collected by the photosystems remains the same (67). Given the phylogenetic distribution of PipX, which is exclusively found in cyanobacteria, it is tempting to propose that it forms part of an early protective response to rapidly decrease photosynthesis upon cold shock.

Finaly, this work calls attention to the importance of performing real time experiments to study the regulation of protein complexes in response to environmentally relevant changes and anticipates further breakthroughs in our understanding of signaling networks.

## MATERIALS AND METHODS

### Plasmid construction

The plasmids and primers used in this study are listed in Table 2 and S2, respectively. *Escherichia coli* XL1-Blue was used to perform Gibson assembly cloning (68). All constructs were verified by automated Sanger sequencing.

The plasmid pUAGC1165 was obtained by assembling fragments F1 and F2, as described by (18). Fragment F1, comprising the *engA* coding region and 162 bp upstream, was amplified by PCR from *S. elongatus* genomic DNA using primers EngA-FL-LgBit-R and SmBiT-EngA-F. Fragment F2 was amplified by PCR from pUAGC1161 using primers FL-LgBiT-4F and SmBiT-2R.

### Cyanobacteria transformation and strain verification

The *S. elongatus* strains used in this study are listed in Table 1. Transformations were performed essentially as described in (69), and the correct insertions were verified by PCR. The primer pairs used were NSI-1R / NS1-2R for NSI, PipX-L80Q-F / LgBit-NS-4R to confirm the presence of *engA*:LgBiT, 2340-For / 2341-Rev for *engA* inactivation, and NS3-seq-1F / NS3-seq-1R for NS3.

### Cyanobacteria growth and culture conditions

*S. elongatus* cultures were routinely grown in blue–green algae BG11 medium (BG11_0_ supplemented with 17.5 mM sodium nitrate (NaNO₃) and 10 mM HEPES/NaOH (pH 7.8); (70)) at 30°C under constant cool white fluorescent light, either in liquid cultures (150 rpm, 70 μmol photons m⁻²s⁻¹; mix of two clones) or on plates (50 μmol photons m⁻²s⁻¹; individual clones).

To modify temperature conditions, cultures grown under standard conditions in flasks were transferred to a Binder KBW 400 or Velp Scientifica™ FOC 2001 Connect incubator, or 500 µL aliquots of the cultures were transferred into 1.5 mL microcentrifuge tubes or 3.5 mL luminometer tubes for each timepoint and incubated in thermostatic water baths.

Solid media contained 1.5% (w/v) agar and, after autoclaving, were supplemented with 0.5 mM sodium thiosulfate (Na₂S₂O₃). The appropriate antibiotic(s) was(were) added at the following concentrations: chloramphenicol (Cm, 3.5 μg/mL), streptomycin (Sm, 15 μg/mL), or nourseothricin (Nt, 15 μg/mL). For liquid growth, cultures of 50 or 170 mL in BG11 were grown in baffled flasks. Growth was monitored by measuring the optical density at 750 nm (OD_750nm_) in 1 mL samples using an Ultrospec 2100 Pro UV-Vis Spectrophotometer (Amersham Biosciences, Amersham, UK). All experiments were performed on mid-exponential phase cultures (OD_750nm_ = 0.4–0.8).

### Protein extraction, immunodetection, and band quantification

For protein extraction, 10 mL samples of cultures were harvested via 8 min centrifugation at 7300× g and stored at −20 °C. The pellets were resuspended in 60 μL of lysis buffer (10 mM Tris/HCl pH 7.5, 0.5 mM EDTA, 1mM β-mercaptoethanol, 1 mM phenylmethylsulfonyl fluoride (PMSF)), and cells were disrupted with 1 spoonful of 0.1 mm glass beads (≈30 µL), as described elsewhere (71). Mixtures were subjected to three cycles of 60 s at a speed of 5 m/s in a high-speed homogenizer Minibeadbeater, followed by 60 s at 4 °C. Samples were centrifuged (5500× g for 5 min), and the supernatant fractions (crude protein extracts) were transferred to a new tube. Protein concentrations were estimated via the Bradford method (72) using the PierceTM detergent-compatible Bradford assay kit (ThermoScientific, Waltham, MA, USA) on a VICTOR3TM 1420 Multilabel Plate Reader. Crude protein extracts were stored at −20°C until needed.

For immunodetection, 10-60 µg of total protein extracts were loaded into a sodium dodecyl sulphate polyacrylamide gel (SDS-PAGE; 15% polyacrylamide). Electrophoresis was followed by immunoblotting onto 0.2 μm polyvinylidene fluoride membranes (from GE Healthcare Technologies, Inc., Chicago, IL, USA), and the membranes were subsequently blocked with Tris-Buffered Saline (TBS-Tween; 20 mM Tris/HCl pH 7.5, 500 mM NaCl, Tween 20 0.1%) solution containing 5% non-fat dried milk for 1 h at room temperature and then incubated overnight in TBS-Tween with 2-5% non-fat dried milk with the corresponding primary antibody. Membranes were then incubated for 1 h at room temperature with a 1:150,000 dilution of ECL rabbit IgG and an HRP-linked F(ab’)2 fragment (from a donkey, GE Healthcare) or a 1:2,500 dilution of mouse IgG (from goat, Merck Millipore, Germany). The signal was detected using a SuperSignal WestFemto reagent (Thermo Fisher Scientific, Waltham, MA, USA) in a Biorad ChemiDoc Imager using the automatic exposure mode and avoiding pixel saturation. A 1:5,000 dilution of primary anti-PipX, anti-EngA, anti-PII, and anti-PlmA antibodies, or a 1:500 (EngA and NtcA) or 1:20,000 (PII) dilution of anti-LgBiT (Promega Corporation) antibody, were used separately.

### NanoBiT Bioluminescence Assays

To measure NanoBiT bioluminescence, 500 µL samples of cyanobacterial cultures were briefly vortexed with 10 µL of a fresh mQ water 13 µM solution of the luciferin Q-108, prepared from Hikarazine-108 as described (73) and incubated for 1 min under the same culture conditions. Bioluminescence was quantified using a luminometer (Junior LB9509, Berthold Technologies GmbH & Co. KG, Bad Wildbad, Germany) with a 5 s measurement time. Raw luminescence values were normalized by the OD_750nm_ of each culture.

### Intracellular ATP Content Determination

The ATP extraction was essentially performed as described in (18). Briefly, 500 µL aliquots were flash-frozen in liquid nitrogen. ATP was extracted via three consecutive cycles of boiling (10 min, 100 °C) and freezing (liquid nitrogen), followed by centrifugation at 14,000× g for 3 min at 4 °C. A 100 µL aliquot of the samples, or an appropriate dilution with mQ water if necessary, was mixed with 40 µL of a reaction solution containing 1 mM DTT, 0.25 mM luciferin, and 7.5 µg/mL luciferase from *Photinus pyralis*. The bioluminescence was measured in black 96-well microplates (OptiPlate-96 F HB; PerkinElmer, Waltham, MA, USA) using a VICTOR3TM 1420 Multilabel Plate Reader (PerkinElmer, Waltham, MA, USA). The ATP content was quantified using the standard curve created in parallel.

### PipX expression and purification

To produce PipX with a N-terminal His_6_ tag (H_6_-PipX), *E. coli* BL21 (DE3) cells, carrying pUAGC73 plasmid, were grown overnight at 37°C with shaking in 50 mL LB-ampicillin (0.1 mg/mL). This preculture was used to inoculate 1 L of LB-ampicillin (0.1 mg/mL) that was grown in the same conditions to an OD_600nm_ of about 0.8. Overexpression was initiated with addition of 1mM isopropyl-β-D-1-thiogalactopyranoside (IPTG), and cells were incubated for 4 h at 30°C. The cells were harvested by centrifugation, washed with 20 mM TrisHCl pH7.4 and 100mM NaCl and frozen at −20°C. The cells were disrupted by sonication in 25 mL of lysis buffer (20mM TrisHCl pH7.4, 400mM NaCl, 50mM KCl, 5mM MgCl_2_, 0.5mM EDTA, 1mM DTT, 1mM benzamidine and 0.2mM phenylmethylsulfonyl fluoride (PMSF)) and the suspension was centrifuged for 20 min at 10,000 x g at 4°C. To purify PipX by Ni-affinity chromatography and imidazole elution, the supernatant was loaded on a 1 mL HisPur™ Ni-NTA Chromatography Cartridge (Thermofisher). The column was washed with 20 mM TrisHCl pH 8.1, 300 mM NaCl and 50 mM imidazole and the bound His_6_-PipX was then eluted with 20 mM TrisHCl pH 8.1, 400 mM NaCl and 500 mM imidazole. The purest fraction (purity checked by SDS-PAGE; 15% polyacrylamide) was concentrated to 0.5 mg/mL (quantified by Bradford method (72) using bovine serum albumin as standard), placing it simultaneously in 20 mM TrisHCl pH 7.4 and 100 mM NaCl by centrifugal ultrafiltration (3 kDa cutoff membrane; Amicon Ultra, from Millipore). Protein was stored at −20°C until use.

### In vitro spectroscopy of isolated recombinant PipX: far-UV circular dichroism and fluorescence

Far-UV circular dichroism: Circular dichroism spectra in the far-ultraviolet region (far-UV CD) were collected on a Jasco J810 (Tokyo, Japan) spectropolarimeter fitted with a thermostated cell holder and interfaced with a Peltier unit. Molar ellipticity was calculated as described previously (74). Isothermal wavelength spectra of PipX at different temperatures (in the range 10°C to 40°C, after equilibration for 10 minutes at the desired temperature), were prepared, acquired and corrected with the same experimental set-up described elsewhere (74, 75). Protein concentration was 12 mM in 50 mM Tris buffer (pH 7.5).

Thermal denaturations of isolated PipX were also carried out by following the changes in the ellipticity at 222 nm as described (75). These experiments allow the determination of the thermal denaturation midpoint of the protein to ensure that the explored temperatures acquired spectra were well below of such value, by using well-known equations (75).

#### Fluorescence

The steady-state spectra of isolated PipX were collected at 25°C on a Cary Varian spectrofluorimeter (Agilent, USA), with a Peltier temperature controller. Sample concentrations and buffer conditions were the same used in the far-UV CD experiments with the same buffer concentration. The experiments were prepared as described elsewhere (74, 75). A 1 cm-pathlength quartz cell (Hellma) was used.

Thermal scans were collected in the same instrument at 302 or 308 nm after excitation at 280 (as PipX has only 8 tyrosine residues, as fluorescence amino acids) from 25 to 85°C with heating rates of 60°C/h and average time of 1 s. The rest of the experimental set-up was the same described above, and additional details have been described elsewhere (74, 75). As it happens with the far-UV CD thermal scans, these experiments allow the determination of the thermal denaturation midpoint of the protein to ensure that the explored temperatures acquired spectra were well below of such value, by using well-known equations (75).

### Computational methods

Protein intensity levels were quantified from Western blot images using *ImageJ* v1.54g. Bands were selected using the “Rectangle” function, and their corresponding intensity profiles were measured with the “Wand” tool. Statistical analyses were performed using *RStudio* (76).

PyMOL (The PyMOL Molecular Graphics System, Version 1.7.1.7, Schrödinger, LLC) was used to generate graphical representations of protein structures.

## Acknowledgments

The authors thank J. L. Yves for kindly providing the hikarazines samples. R. Cantos, C. Jerez, and P. Salinas for technical contributions or advice. This work was supported by grant PID2023-149456NB-I00, funded by MCIN/AEI/10.13039/501100011033 from the Spanish Government, and grants VIGROB23-126 and GRE20-04-C from the University of Alicante to A.C and by Horizon European Union [EXPLORA GA n° 101181841] to JLN. R.D. was supported by the UKRI-BBSRC (grant BBS/E/J/000PR9797) and by the Royal Society (grant ICA\R1\180088). L.F-G. was the recipient of a Grant for official master’s degree studies and initiation to research from the Office of the Vice President of Research of the University of Alicante. S.B. was supported by a National Grant from the Algerian Ministry of Higher Education and Scientific Research.

## SUPPLEMENTAL MATERIAL(S)

**Figure S1.**
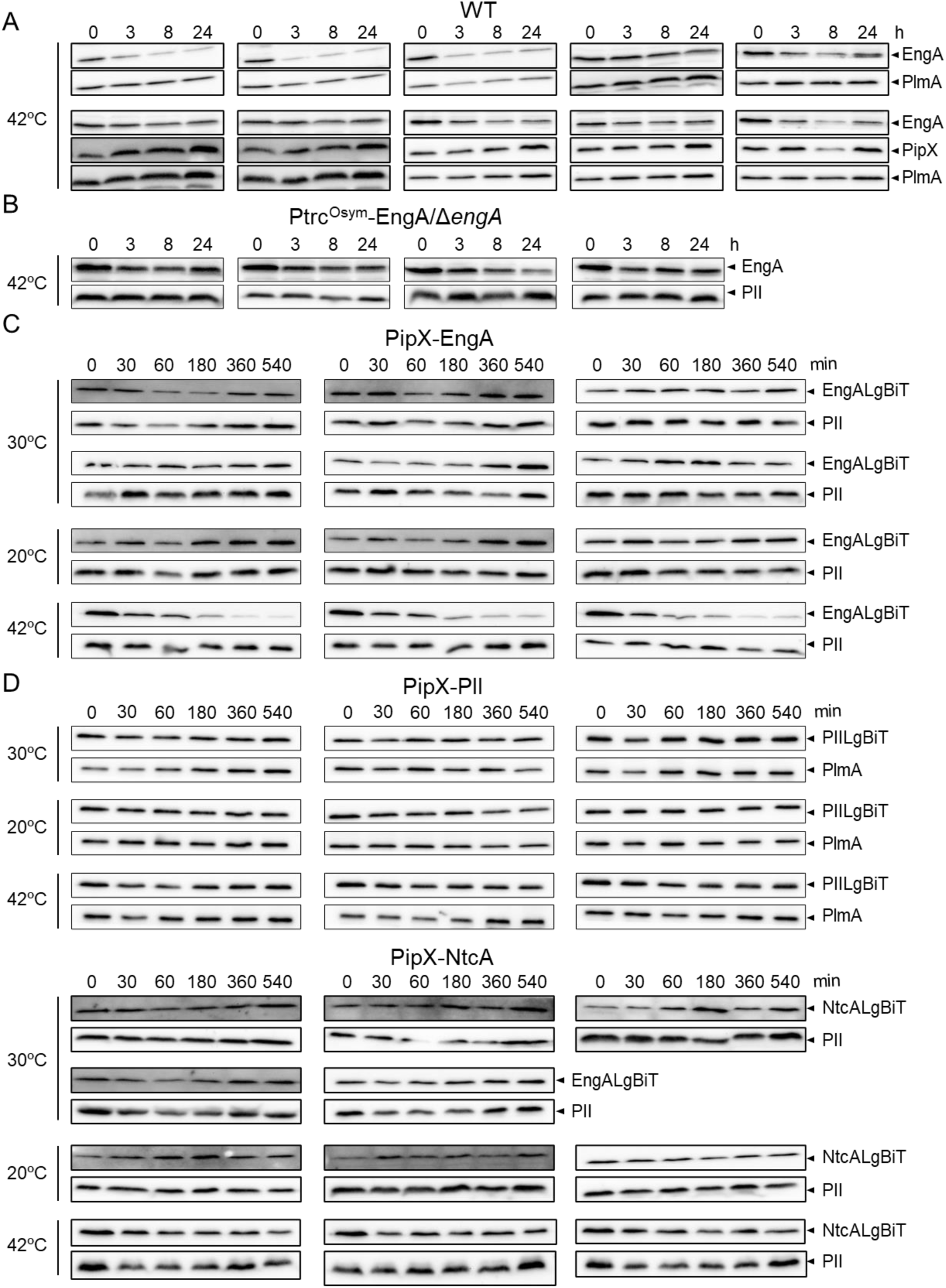
Immunodetection analysis of the indicated proteins for the quantifications performed in Fig. 2A (*A*), Fig. 2B (*B*), Fig. 3D (*C*), and Fig. 4B (*D*).

**Figure S2.**
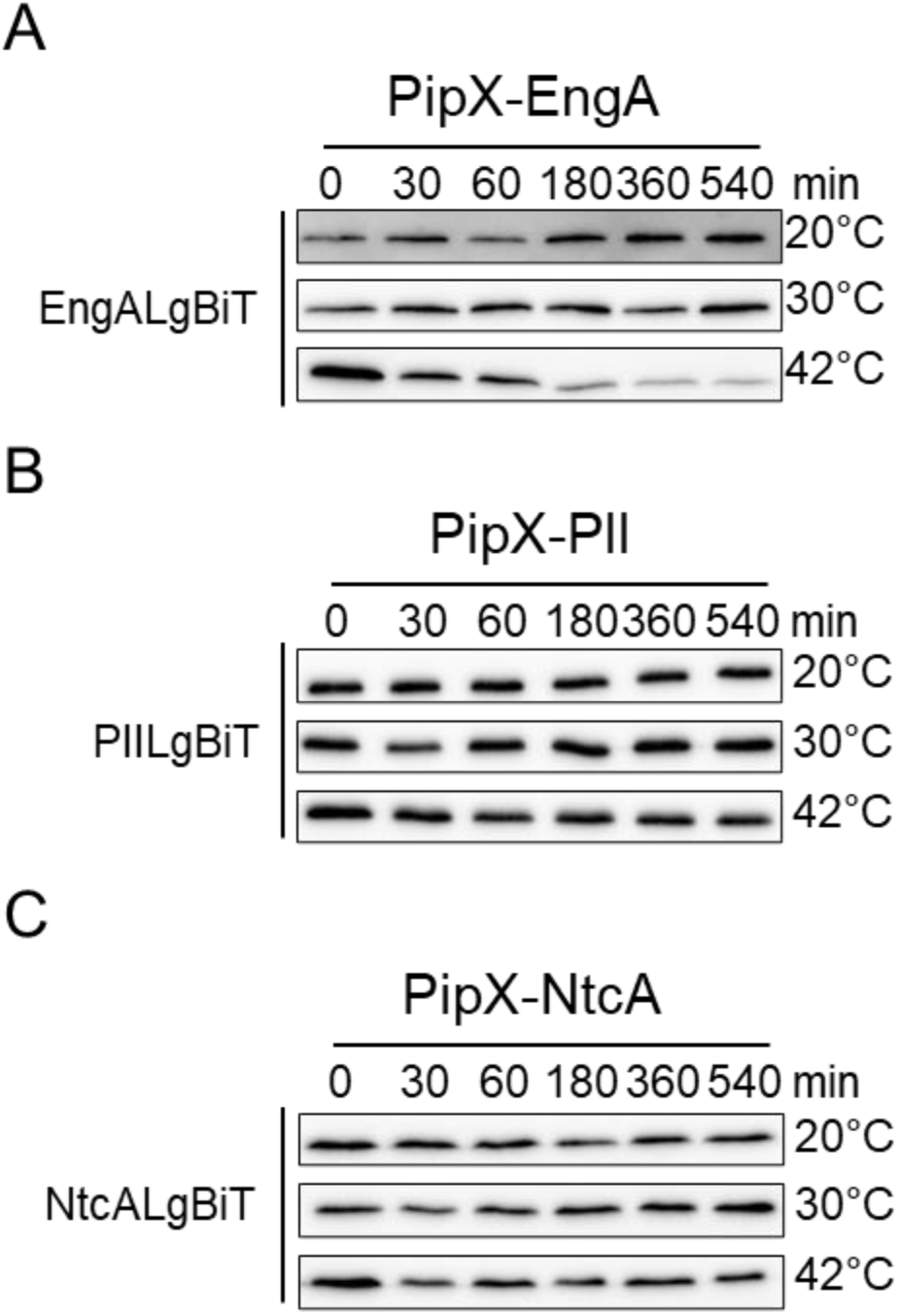
Representative immunodetection of the indicated proteins and conditions for the quantifications performed in Fig. 3D (*A*) and Fig. 4B (*B, C*).

**Figure S3.**
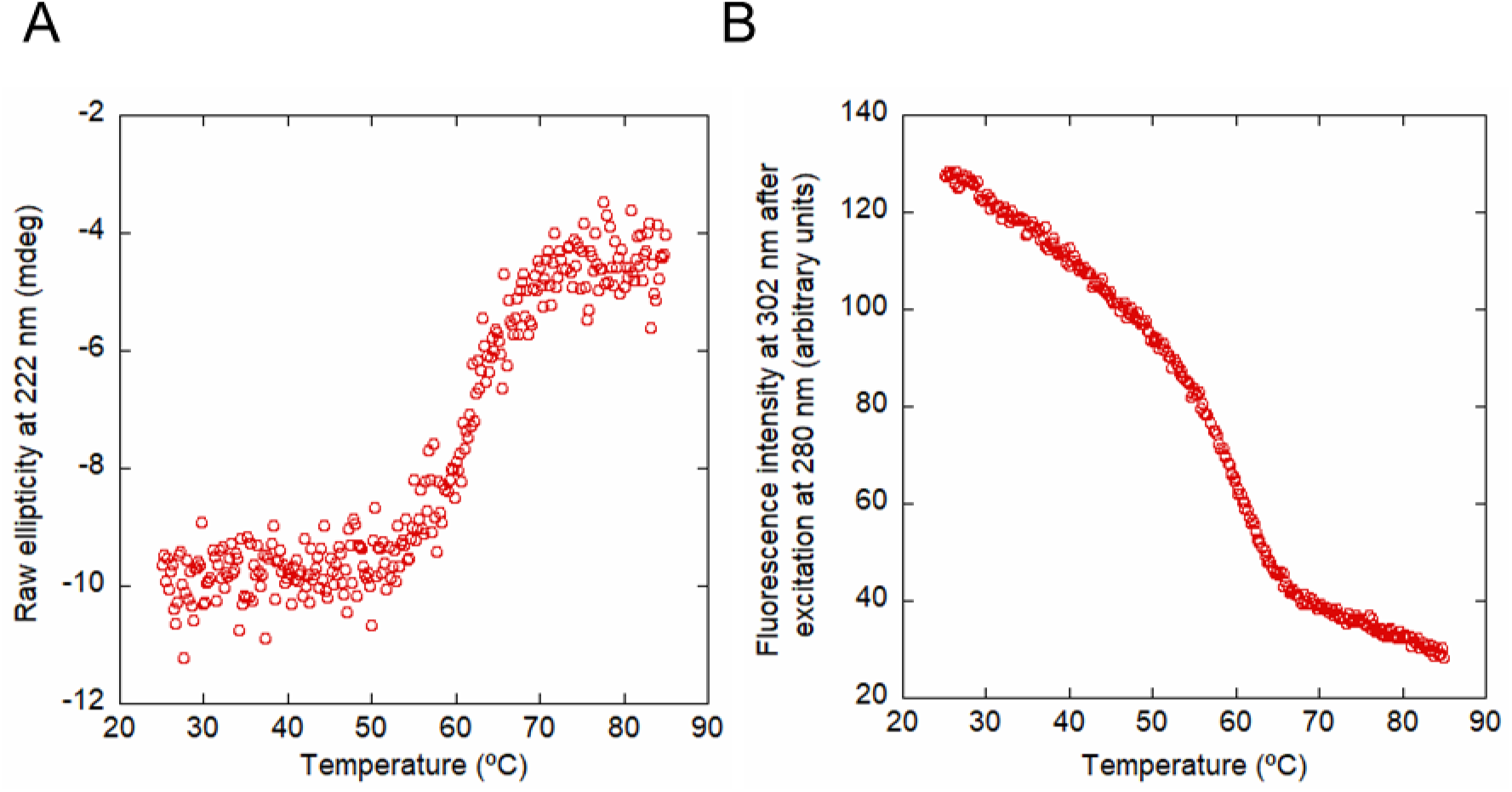
Thermal denaturation of PipX. Thermal denaturation of isolated recombinant PipX followed by the ellipticity and 222 nm in 50 mM Tris (pH 7.5) (*A*), and by the intrinsic fluorescence emission at 302 nm after excitation at 280 nm in 50 mM Tris (pH 7.5).

**Table S1.** ATP levels in response to temperature up or downshifts. Relative ATP levels, normalized with the OD_750_ and referred to timepoint 0, in cultures incubated at 20°C, 30°C, or 42°C for the indicated times. Data are presented as means with error bars (±standard deviation) from at least six (0-60 min) or three (180-540 min) biological replicates. Welch’s t-test with Bonferroni correction was used to compare data between 30°C and either 20°C or 42°C at the same timepoint. Significant differences (p ≤ 0.05 (*)) were detected at the 5-min timepoint for 20°C.

| Temperature | Time (min) |  |  |  |  |  |  |  |  |
| --- | --- | --- | --- | --- | --- | --- | --- | --- | --- |
|  | 0 | 2 | 5 | 15 | 30 | 60 | 180 | 360 | 540 |
| 20°C | 1 | 0.88<br>$\pm 0.07$ | 0.72*<br>$\pm 0.08$ | 0.76<br>$\pm 0.16$ | 0.82<br>$\pm 0.14$ | 0.80<br>$\pm 0.20$ | 0.72<br>$\pm 0.12$ | 0.73<br>$\pm 0.11$ | 0.72<br>$\pm 0.04$ |
| 30°C | 1 | 0.96<br>$\pm 0.09$ | 0.89<br>$\pm 0.12$ | 0.90<br>$\pm 0.11$ | 0.90<br>$\pm 0.10$ | 0.87<br>$\pm 0.08$ | 0.89<br>$\pm 0.07$ | 0.76<br>$\pm 0.09$ | 0.91<br>$\pm 0.29$ |
| 42°C | 1 | 0.94<br>$\pm 0.19$ | 1.00<br>$\pm 0.23$ | 1.05<br>$\pm 0.28$ | 0.93<br>$\pm 0.19$ | 0.83<br>$\pm 0.24$ | 0.81<br>$\pm 0.12$ | 0.87<br>$\pm 0.02$ | 0.81<br>$\pm 0.10$ |

**Table S2.**
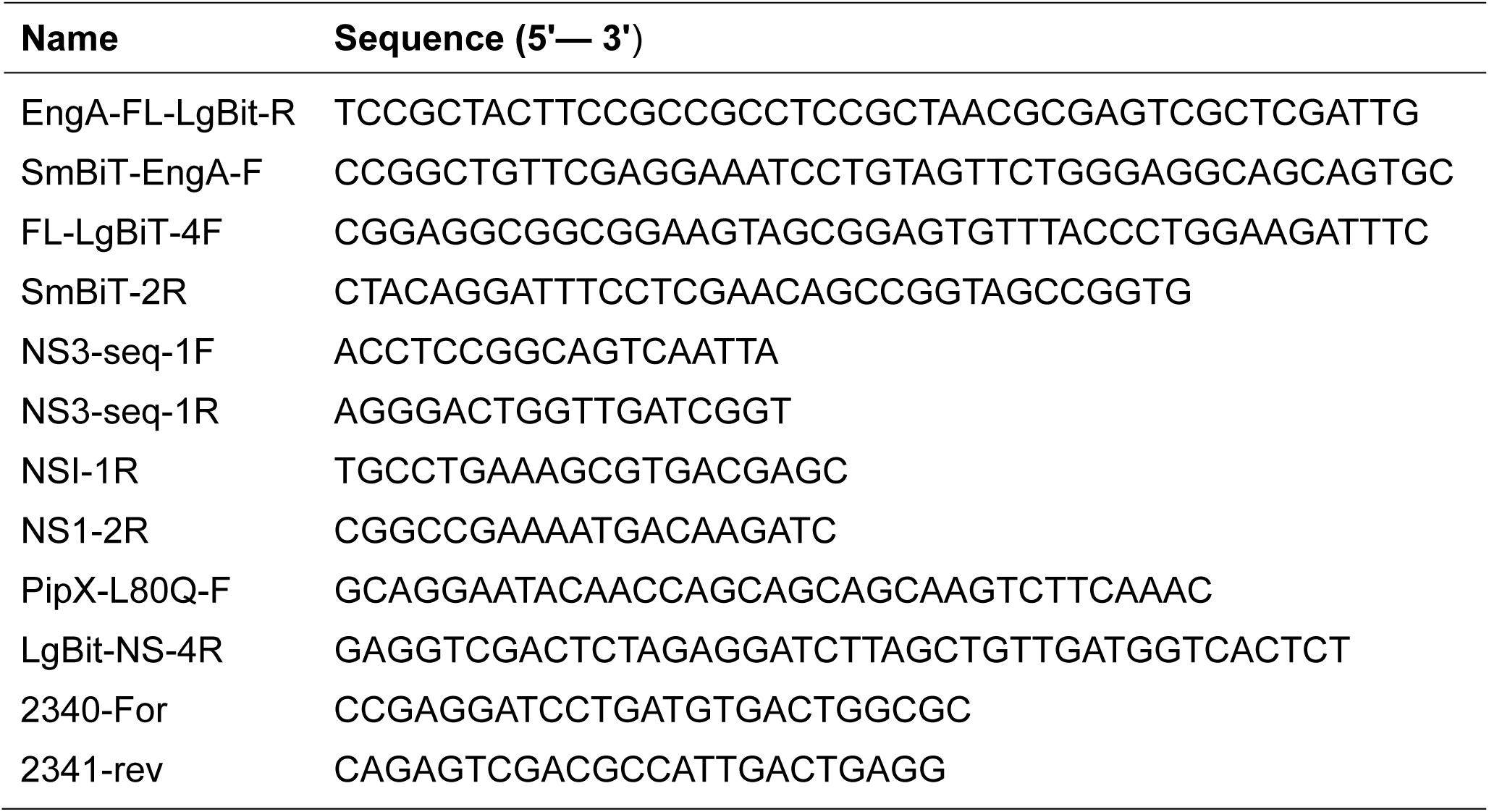
Oligonucleotides.

## REFERENCES

1. Blank CE, Sánchez-Baracaldo P. 2010. Timing of morphological and ecological innovations in the cyanobacteria – a key to understanding the rise in atmospheric oxygen. Geobiology 8:1–23.

2. Lee H-W, Noh J-H, Choi D-H, Yun M, Bhavya PS, Kang J-J, Lee J-H, Kim K-W, Jang H-K, Lee S-H. 2021. Picocyanobacterial Contribution to the Total Primary Production in the Northwestern Pacific Ocean. Water (Basel) 13:1610.

3. Khan S, Fu P. 2020. Biotechnological perspectives on algae: a viable option for next generation biofuels. Curr Opin Biotechnol 62:146–152.

4. Zhang C-C, Zhou C-Z, Burnap RL, Peng L. 2018. Carbon/Nitrogen Metabolic Balance: Lessons from Cyanobacteria. Trends Plant Sci 23:1116–1130.

5. Forchhammer K, Selim KA. 2020. Carbon/nitrogen homeostasis control in cyanobacteria. FEMS Microbiol Rev 44:33–53.

6. Forchhammer K, Selim KA, Huergo LF. 2022. New views on PII signaling: from nitrogen sensing to global metabolic control. Trends Microbiol 30:722–735.

7. Zeth K, Fokina O, Forchhammer K. 2014. Structural Basis and Target-specific Modulation of ADP Sensing by the Synechococcus elongatus PII Signaling Protein. Journal of Biological Chemistry 289:8960–8972.

8. Kamberov ES, Atkinson MR, Ninfa AJ. 1995. The Escherichia coli PII Signal Transduction Protein Is Activated upon Binding 2-Ketoglutarate and ATP. Journal of Biological Chemistry 270:17797–17807.

9. Espinosa J, Forchhammer K, Burillo S, Contreras A. 2006. Interaction network in cyanobacterial nitrogen regulation: PipX, a protein that interacts in a 2-oxoglutarate dependent manner with PII and NtcA. Mol Microbiol 61:457–469.

10. Herrero A, Muro-Pastor AM, Flores E. 2001. Nitrogen Control in Cyanobacteria. J Bacteriol 183:411–425.

11. Esteves-Ferreira AA, Inaba M, Fort A, Araújo WL, Sulpice R. 2018. Nitrogen metabolism in cyanobacteria: metabolic and molecular control, growth consequences and biotechnological applications. Crit Rev Microbiol 44:541–560.

12. Labella JI, Cantos R, Salinas P, Espinosa J, Contreras A. 2020. Distinctive Features of PipX, a Unique Signaling Protein of Cyanobacteria. Life 10:79.

13. Burillo S, Luque I, Fuentes I, Contreras A. 2004. Interactions between the Nitrogen Signal Transduction Protein PII and *N* -Acetyl Glutamate Kinase in Organisms That Perform Oxygenic Photosynthesis. J Bacteriol 186:3346–3354.

14. Espinosa J, Labella JI, Cantos R, Contreras A. 2018. Energy drives the dynamic localization of cyanobacterial nitrogen regulators during diurnal cycles. Environ Microbiol 20:1240–1252.

15. Espinosa J, Forchhammer K, Contreras A. 2007. Role of the Synechococcus PCC 7942 nitrogen regulator protein PipX in NtcA-controlled processes. Microbiology (N Y) 153:711–718.

16. Llácer JL, Espinosa J, Castells MA, Contreras A, Forchhammer K, Rubio V. 2010. Structural basis for the regulation of NtcA-dependent transcription by proteins PipX and PII. Proceedings of the National Academy of Sciences 107:15397–15402.

17. Laichoubi KB, Beez S, Espinosa J, Forchhammer K, Contreras A. 2011. The nitrogen interaction network in Synechococcus WH5701, a cyanobacterium with two PipX and two PII-like proteins. Microbiology (N Y) 157:1220–1228.

18. Jerez C, Llop A, Salinas P, Bibak S, Forchhammer K, Contreras A. 2024. Analysing the Cyanobacterial PipX Interaction Network Using NanoBiT Complementation in Synechococcus elongatus PCC7942. Int J Mol Sci 25:4702.

19. Llop A, Tremiño L, Cantos R, Contreras A. 2023. The Signal Transduction Protein PII Controls the Levels of the Cyanobacterial Protein PipX. Microorganisms 11:2379.

20. Zhao M-X, Jiang Y-L, Xu B-Y, Chen Y, Zhang C-C, Zhou C-Z. 2010. Crystal Structure of the Cyanobacterial Signal Transduction Protein PII in Complex with PipX. J Mol Biol 402:552–559.

21. Zhao M-X, Jiang Y-L, He Y-X, Chen Y-F, Teng Y-B, Chen Y, Zhang C-C, Zhou C-Z. 2010. Structural basis for the allosteric control of the global transcription factor NtcA by the nitrogen starvation signal 2-oxoglutarate. Proceedings of the National Academy of Sciences 107:12487–12492.

22. Espinosa J, Castells MA, Laichoubi KB, Contreras A. 2009. Mutations at *pipX* Suppress Lethality of P _II_ -Deficient Mutants of *Synechococcus elongatus* PCC 7942. J Bacteriol 191:4863–4869.

23. Espinosa J, Castells MA, Laichoubi KB, Forchhammer K, Contreras A. 2010. Effects of spontaneous mutations in PipX functions and regulatory complexes on the cyanobacterium Synechococcus elongatus strain PCC 7942. Microbiology (N Y) 156:1517–1526.

24. Espinosa J, Rodríguez-Mateos F, Salinas P, Lanza VF, Dixon R, de la Cruz F, Contreras A. 2014. PipX, the coactivator of NtcA, is a global regulator in cyanobacteria. Proceedings of the National Academy of Sciences 111:201404030–201404097.

25. Forcada-Nadal A, Llácer JL, Contreras A, Marco-Marín C, Rubio V. 2018. The PII-NAGK-PipX-NtcA Regulatory Axis of Cyanobacteria: A Tale of Changing Partners, Allosteric Effectors and Non-covalent Interactions. Front Mol Biosci 5:91.

26. Laichoubi KB, Espinosa J, Castells MA, Contreras A. 2012. Mutational Analysis of the Cyanobacterial Nitrogen Regulator PipX. PLoS One 7:e35845.

27. Forcada-Nadal A, Bibak S, Salinas P, Contreras A, Rubio V, Llácer JL. 2025. Structures of the cyanobacterial nitrogen regulators NtcA and PipX complexed to DNA shed light on DNA binding by NtcA and implicate PipX in the recruitment of RNA polymerase. Nucleic Acids Res 53.

28. Giner-Lamia J, Robles-Rengel R, Hernández-Prieto MA, Muro-Pastor MI, Florencio FJ, Futschik ME. 2017. Identification of the direct regulon of NtcA during early acclimation to nitrogen starvation in the cyanobacterium Synechocystis sp. PCC 6803. Nucleic Acids Res 45:11800–11820.

29. Jerez C, Salinas P, Llop A, Cantos R, Espinosa J, Labella JI, Contreras A. 2021. Regulatory Connections Between the Cyanobacterial Factor PipX and the Ribosome Assembly GTPase EngA. Front Microbiol 12:781760–NA.

30. Llop A, Bibak S, Cantos R, Salinas P, Contreras A. 2023. The ribosome assembly GTPase EngA is involved in redox signaling in cyanobacteria. Front Microbiol 14:1242616–NA.

31. Sinetova MA, Los DA. 2016. New insights in cyanobacterial cold stress responses: Genes, sensors, and molecular triggers. Biochimica et Biophysica Acta (BBA) - General Subjects 1860:2391–2403.

32. Kobayashi I, Watanabe S, Kanesaki Y, Shimada T, Yoshikawa H, Tanaka K. 2017. Conserved two-component <scp>H</scp> ik34-<scp>R</scp> re1 module directly activates heat-stress inducible transcription of major chaperone and other genes in *Synechococcus elongatus* PCC7942. Mol Microbiol 104:260–277.

33. Mironov K, Sinetova M, Shumskaya M, Los D. 2019. Universal Molecular Triggers of Stress Responses in Cyanobacterium Synechocystis. Life 9:67.

34. Weber MHW, Marahiel MA. 2003. Bacterial Cold Shock Responses. Sci Prog 86:9–75.

35. Shivaji S, Prakash JSS. 2010. How do bacteria sense and respond to low temperature? Arch Microbiol 192:85–95.

36. Zhang Y, Gross CA. 2021. Cold Shock Response in Bacteria. Annu Rev Genet 55:377–400.

37. Kusukawa N, Yura T. 1988. Heat shock protein GroE of Escherichia coli: key protective roles against thermal stress. Genes Dev 2:874–882.

38. Yura T. 2019. Regulation of the heat shock response in &lt;i&gt;Escherichia coli&lt;/i&gt;: history and perspectives. Genes Genet Syst 94:103–108.

39. Schumann W. 2016. Regulation of bacterial heat shock stimulons. Cell Stress Chaperones 21:959–968.

40. Dixon AS, Schwinn MK, Hall MP, Zimmerman K, Otto P, Lubben TH, Butler BL, Binkowski BF, Machleidt T, Kirkland TA, Wood MG, Eggers CT, Encell LP, Wood K V. 2016. NanoLuc Complementation Reporter Optimized for Accurate Measurement of Protein Interactions in Cells. ACS Chem Biol 11:400–408.

41. Kashima D, Kageoka M, Kimura Y, Horikawa M, Miura M, Nakakido M, Tsumoto K, Nagamune T, Kawahara M. 2021. A Novel Cell-Based Intracellular Protein–Protein Interaction Detection Platform (SOLIS) for Multimodality Screening. ACS Synth Biol 10:990–999.

42. Pipchuk A, Yang X. 2021. Using Biosensors to Study Protein–Protein Interaction in the Hippo Pathway. Front Cell Dev Biol 9.

43. Sicking M, Jung M, Lang S. 2021. Lights, Camera, Interaction: Studying Protein–Protein Interactions of the ER Protein Translocase in Living Cells. Int J Mol Sci 22:10358.

44. Bardelang P, Murray EJ, Blower I, Zandomeneghi S, Goode A, Hussain R, Kumari D, Siligardi G, Inoue K, Luckett J, Doutch J, Emsley J, Chan WC, Hill P, Williams P, Bonev BB. 2023. Conformational analysis and interaction of the Staphylococcus aureus transmembrane peptidase AgrB with its AgrD propeptide substrate. Front Chem 11.

45. Oliveira Paiva AM, Friggen AH, Qin L, Douwes R, Dame RT, Smits WK. 2019. The Bacterial Chromatin Protein HupA Can Remodel DNA and Associates with the Nucleoid in Clostridium difficile. J Mol Biol 431:653–672.

46. Rozbeh R, Forchhammer K. 2024. In Vivo Detection of Metabolic Fluctuations in Real Time Using the NanoBiT Technology Based on PII Signalling Protein Interactions. Int J Mol Sci 25:3409.

47. Rozbeh R, Forchhammer K. 2021. Split NanoLuc technology allows quantitation of interactions between PII protein and its receptors with unprecedented sensitivity and reveals transient interactions. Sci Rep 11:12535.

48. Westerhausen S, Nowak M, Torres-Vargas CE, Bilitewski U, Bohn E, Grin I, Wagner S. 2020. A NanoLuc luciferase-based assay enabling the real-time analysis of protein secretion and injection by bacterial type III secretion systems. Mol Microbiol 113:1240–1254.

49. Bharat A, Brown ED. 2014. Phenotypic investigations of the depletion of EngA in *Escherichia coli* are consistent with a role in ribosome biogenesis. FEMS Microbiol Lett 353:26–32.

50. Kelly SM, Price NC. 2006. Circular Dichroism to Study Protein Interactions. Curr Protoc Protein Sci 46.

51. Forcada-Nadal A, Palomino-Schätzlein M, Neira JL, Pineda-Lucena A, Rubio V. 2017. The PipX Protein, When Not Bound to Its Targets, Has Its Signaling C-Terminal Helix in a Flexed Conformation. Biochemistry 56:3211–3224.

52. Los DA, Zorina A, Sinetova M, Kryazhov S, Mironov K, Zinchenko V V. 2010. Stress Sensors and Signal Transducers in Cyanobacteria. Sensors 10:2386– 2415.

53. Selim KA, Alva V. 2024. PII-like signaling proteins: a new paradigm in orchestrating cellular homeostasis. Curr Opin Microbiol 79:102453.

54. Gupta KR, Baloni P, Indi SS, Chatterji D. 2016. Regulation of Growth, Cell Shape, Cell Division, and Gene Expression by Second Messengers (p)ppGpp and Cyclic Di-GMP in Mycobacterium smegmatis. J Bacteriol 198:1414–1422.

55. Boniecka J, Prusińska J, Dąbrowska GB, Goc A. 2017. Within and beyond the stringent response-RSH and (p)ppGpp in plants. Planta 246:817–842.

56. Corrigan RM, Bellows LE, Wood A, Gründling A. 2016. ppGpp negatively impacts ribosome assembly affecting growth and antimicrobial tolerance in Gram-positive bacteria. Proceedings of the National Academy of Sciences 113.

57. Zhang Y, Burkhardt DH, Rouskin S, Li G-W, Weissman JS, Gross CA. 2018. A Stress Response that Monitors and Regulates mRNA Structure Is Central to Cold Shock Adaptation. Mol Cell 70:274–286.e7.

58. Mehrez M, Romand S, Field B. 2023. New perspectives on the molecular mechanisms of stress signalling by the nucleotide guanosine tetraphosphate (ppGpp), an emerging regulator of photosynthesis in plants and algae. New Phytologist 237:1086–1099.

59. Wang B, Dai P, Ding D, Del Rosario A, Grant RA, Pentelute BL, Laub MT. 2019. Affinity-based capture and identification of protein effectors of the growth regulator ppGpp. Nat Chem Biol 15:141–150.

60. Hood RD, Higgins SA, Flamholz A, Nichols RJ, Savage DF. 2016. The stringent response regulates adaptation to darkness in the cyanobacterium *Synechococcus elongatus*. Proceedings of the National Academy of Sciences 113:201524915–201524976.

61. Puszynska AM, O’Shea EK. 2017. ppGpp Controls Global Gene Expression in Light and in Darkness in S. elongatus. Cell Rep 21:3155–3165.

62. Llop A, Labella JI, Borisova M, Forchhammer K, Selim KA, Contreras A. 2023. Pleiotropic effects of PipX, PipY, or RelQ overexpression on growth, cell size, photosynthesis, and polyphosphate accumulation in the cyanobacterium Synechococcus elongatus PCC7942. Front Microbiol 14:1141775–NA.

63. Labella JI, Cantos R, Espinosa J, Forcada-Nadal A, Rubio V, Contreras A. 2017. PipY, a Member of the Conserved COG0325 Family of PLP-Binding Proteins, Expands the Cyanobacterial Nitrogen Regulatory Network. Front Microbiol 8:1244.

64. Riediger M, Spät P, Bilger R, Voigt K, Maček B, Hess WR. 2021. Analysis of a photosynthetic cyanobacterium rich in internal membrane systems via gradient profiling by sequencing (Grad-seq). Plant Cell 33:248–269.

65. Jansz ER, Maclean FI. 1973. The effect of cold shock on the blue-green alga *Anacystis nidulans*. Can J Microbiol 19:381–387.

66. Mackey KRM, Paytan A, Caldeira K, Grossman AR, Moran D, McIlvin M, Saito MA. 2013. Effect of Temperature on Photosynthesis and Growth in Marine Synechococcus spp. Plant Physiol 163:815–829.

67. Imlay JA. 2003. Pathways of Oxidative Damage. Annu Rev Microbiol 57:395– 418.

68. Gibson DG, Young L, Chuang R-Y, Venter JC, Hutchison CA, Smith HO. 2009. Enzymatic assembly of DNA molecules up to several hundred kilobases. Nat Methods 6:343–345.

69. Taton A, Erikson C, Yang Y, Rubin BE, Rifkin SA, Golden JW, Golden SS. 2020. The circadian clock and darkness control natural competence in cyanobacteria. Nat Commun 11:1688.

70. Rippka R, Deruelles J, Waterbury JB, Herdman M, Stanier RY. 1979. Generic Assignments, Strain Histories and Properties of Pure Cultures of Cyanobacteria. Microbiology (N Y) 111:1–61.

71. Labella JI, Obrebska A, Espinosa J, Salinas P, Forcada-Nadal A, Tremiño L, Rubio V, Contreras A. 2016. Expanding the Cyanobacterial Nitrogen Regulatory Network: The GntR-Like Regulator PlmA Interacts with the PII-PipX Complex. Front Microbiol 7:1677.

72. Bradford M. 1976. A Rapid and Sensitive Method for the Quantitation of Microgram Quantities of Protein Utilizing the Principle of Protein-Dye Binding. Anal Biochem 72:248–254.

73. Coutant EP, Gagnot G, Hervin V, Baatallah R, Goyard S, Jacob Y, Rose T, Janin YL. 2020. Bioluminescence Profiling of NanoKAZ/NanoLuc Luciferase Using a Chemical Library of Coelenterazine Analogues. Chemistry – A European Journal 26:948–958.

74. Neira JL, Román-Trufero M, Contreras LM, Prieto J, Singh G, Barrera FN, Renart ML, Vidal M. 2009. The Transcriptional Repressor RYBP Is a Natively Unfolded Protein Which Folds upon Binding to DNA. Biochemistry 48:1348– 1360.

75. Czypionka A, Ruiz de los Paños O, Mateu MG, Barrera FN, Hurtado-Gómez E, Gómez J, Vidal M, Neira JL. 2007. The Isolated C-Terminal Domain of Ring1B Is a Dimer Made of Stable, Well-Structured Monomers. Biochemistry 46:12764– 12776.

76. RStudio: Integrated Development for R. RStudio. 2020. 2023.12.1 Build 402. RStudio, PBC, Boston, MA, USA.

77. Eswaramoorthy S, Gerchman S, Graziano V, Kycia H, Studier FW, Swaminathan S. 2003. Structure of a yeast hypothetical protein selected by a structural genomics approach. Acta Crystallogr D Biol Crystallogr 59:127–135.

78. Bullock WO, Fernandez JM, Short JM. 1987. XL1-Blue—a high-efficiency plasmid transforming recA Escherichia coli strain with β-galactosidase selection. Biotechniques 5:376–379.

